# Amplification-free Detection of Zoonotic Viruses Using Cas13 and Multiple CRISPR RNAs

**DOI:** 10.1101/2025.05.15.654407

**Authors:** Caitlin H. Lamb, Aartjan J. W. te Velthuis, Cameron Myhrvold, Benjamin E. Nilsson-Payant

**Affiliations:** Department of Molecular Biology, Princeton University, Princeton, NJ 08544; Department of Chemical and Biological Engineering, Princeton University, Princeton, NJ 08544; Omenn-Darling Bioengineering Institute, Princeton University, Princeton, NJ 08544; Department of Chemistry, Princeton University, Princeton, NJ 08544; Institute for Experimental Virology, TWINCORE, Centre for Experimental and Clinical Infection Research, 30625 Hannover, Germany; Cluster of Excellence RESIST (EXC2155), Hannover Medical School, 30625 Hannover, Germany; Department of Microbiology, Tumor and Cell Biology, Karolinska Institutet, 17165 Stockholm, Sweden

## Abstract

Zoonotic viruses such as hantaviruses and influenza A viruses present a threat to humans and livestock. There is thus a need for methods that are rapid, sensitive, and relatively cheap to detect infections with these pathogens early. Here we use an amplification-free CRISPR-Cas13-based assay, which is simple, cheap and field-deployable, to detect the presence or absence of genomic hantavirus or influenza A virus RNA. In addition, we evaluate whether the use of multiple CRISPR RNAs (crRNAs) can improve the sensitivity of this amplification-free method. We demonstrate that for the hantaviruses Tula Virus (TULV) and Andes Virus (ANDV) a combination of two or three crRNAs provides the best sensitivity for detecting viral RNA, whereas for influenza virus RNA detection, additional crRNAs provide no benefit. We also show that the amplification-free method can be used to detect TULV and ANDV RNA in tissue culture infection samples and influenza A virus RNA in clinical nasopharyngeal swabs. In clinical samples, the Cas13 assay has an 85% agreement with RT-qPCR for identifying a positive sample. Overall, these findings indicate that amplification-free CRISPR-Cas13 detection of viral RNA has potential as a tool for rapidly detecting zoonotic virus infections.

## Introduction

Zoonotic viruses, such as hantaviruses and influenza viruses, pose serious threats to human health. Hantaviruses are a family of tri-segmented negative-sense RNA viruses that naturally circulate in rodents and other small mammals, where they cause life-long asymptomatic infections (1, 2). Viral particles are shed in the urine and feces of infected animals, and zoonotic infections of humans can occur by the inhalation of dried or aerosolized secretions. Clinical manifestations of hantavirus infections in humans can range from asymptomatic or mild flu-like symptoms to highly pathogenic diseases, including hemorrhagic fever with renal syndrome (HFRS) and hantavirus cardiopulmonary syndrome (HCPS) with mortality rates of up to 50%. Annually up to 200,000 human hantavirus cases are recorded, however, due to the largely asymptomatic or mild flu-like symptoms in most human infections, it is likely that actual case numbers are significantly higher (3). While humans are typically considered to be dead-end hosts for most hantaviruses, exceptions exist. For instance, the highly pathogenic Andes virus (ANDV) can transmit between humans by an airborne route. Coupled to their pathogenicity and the ubiquitous presence of rodents and small mammals around human populations, this potential for airborne human-to-human transmission makes hantaviruses like ANDV a significant pandemic and global health threat.

Influenza A viruses (IAV) are also single-stranded negative-sense viruses with a segmented genome (4, 5). IAV strains cause seasonal epidemics and occasional pandemics, with disease typically manifesting as a mild to severe-respiratory illness. IAV has a wide range of mammalian and avian hosts, such as birds, pigs, cows, and humans, which creates many opportunities for zoonotic spillover (5). Moreover, co-infection of a single host with multiple IAV strains can lead to reassortment, producing genetically novel viruses with pandemic potential. A recent example is the 2009 H1N1 IAV pandemic, which was caused by a spillover event from a pig infected with a triple-reassorted IAV strain (5). Thus, as with hantaviruses, animal reservoirs play a critical role in the emergence of new IAV strains.

To address the looming threat posed by emerging zoonotic RNA viruses, we rely on fast detection methods for surveillance and early diagnosis. Several molecular tools are available to diagnose and study influenza viruses, including RT-qPCR, primer extension, sequencing, and antigen-based techniques. However, the tools to diagnose and study hantaviruses are more limited. Currently, hantavirus detection relies on serology for IgG and IgM, and RT-qPCR (6). Serology and RT-qPCR are both labor- and time-intensive and require equipment and laboratory infrastructure to be conducted. Given these challenges, additional strategies are needed to detect hantaviruses and emerging IAVs.

One promising strategy for the surveillance of zoonotic viruses is CRISPR-Cas13-based nucleic acid detection (7–10). CRISPR-Cas13-based detection methods harness the *trans*-cleavage activity of the RNA-guided RNase, Cas13, along with an RNA reporter, allowing for enhanced sensitivity and specificity relative to hybridization alone (7, 8, 11, 12). Importantly, these methods can be used to directly detect viral RNA (vRNA) without the need for reverse transcription or nucleic acid amplification (13–15). Furthermore, it has been shown that using multiple CRISPR RNAs (crRNAs) to target different sequences in the same RNA molecule can increase the sensitivity of SARS-CoV-2 detection (Figure 1) (13).

**Figure 1.**
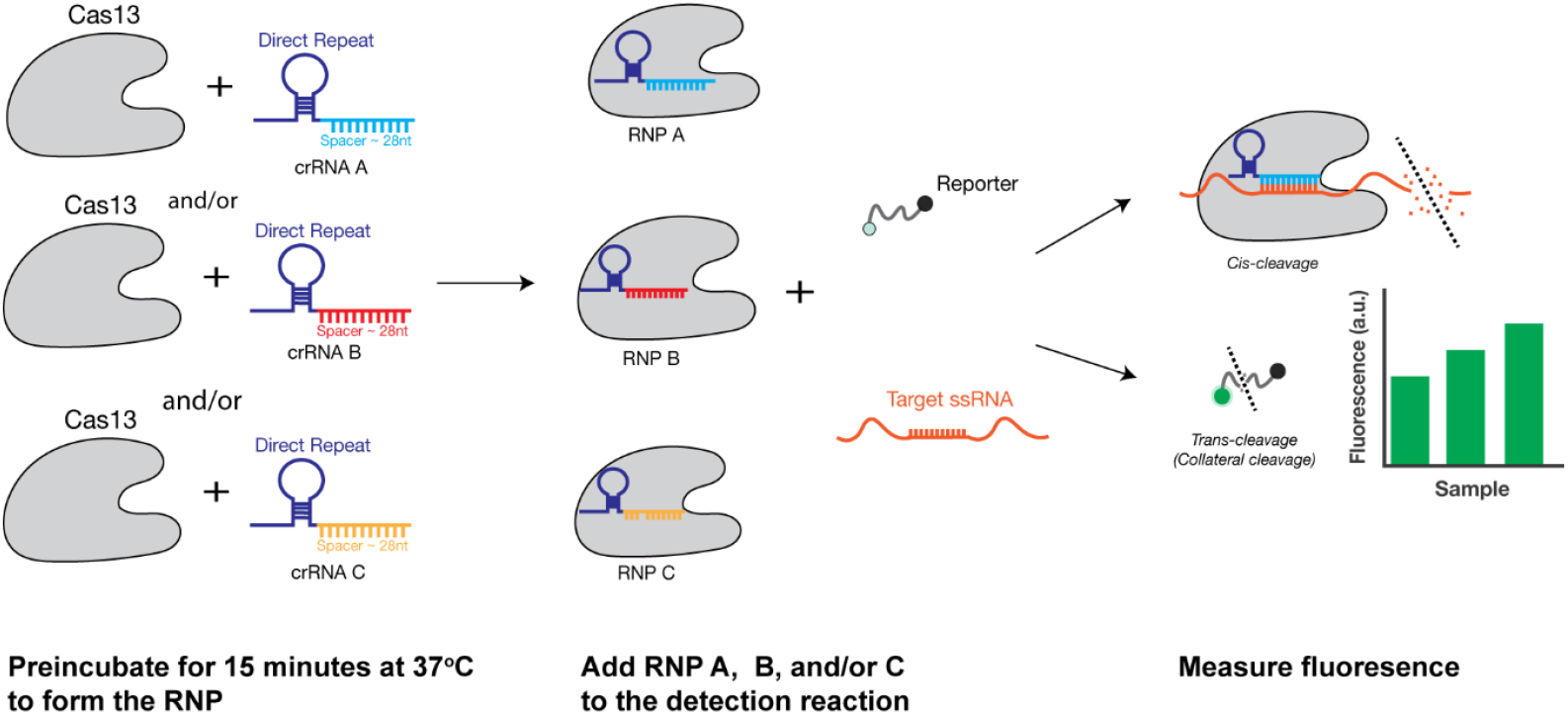
Workflow of multi-crRNA Cas13-based amplification-free detection. Cas13 is first complexed to the crRNA to form the ribonucleoprotein complex ((RNP) A, B, or C). The RNP is then added to the detection reaction containing the fluorescent reporter and target RNA. The Cas13 RNP finds the target RNA, activates, and cleaves the reporter in *trans*, generating a fluorescent signal.

Although Cas13 assays that detect hantavirus and IAV RNA using amplification have been reported (10, 16), an amplification-free assay would be simpler and easier for rapid screening of suspicious samples. We previously used an amplification-free Cas13-based method to detect aberrant IAV RNAs (15). Here, we sought to use amplification-free CRISPR-Cas13 to detect two hantaviruses, Tula virus (TULV) and ANDV, in tissue culture infection samples using one or multiple crRNAs. In addition, we explored whether this method can be applied to the detection of IAV genomic RNA. We found that using multiple crRNAs enhances detection sensitivity for both the highly pathogenic New World hantavirus ANDV and the mostly non-pathogenic Old World hantavirus TULV. However, in the case of IAV, multiple crRNAs did not improve the detection sensitivity. We therefore conclude that amplification-free Cas13 detection using multiple crRNAs has the potential to be a powerful approach to detect zoonotic virus infection, but that it does not always increase the sensitivity of the Cas13 detection and that the assay will need to be optimized for each virus separately.

## Results

### Combining crRNAs can enhance hantavirus detection

Currently, there are limited tools and methods to detect and study hantaviruses. As previously shown for IAV, RT-PCR or RT-qPCR may introduce sequence specific amplification biases (15). In addition, these methods require the end user to optimize temperature, primer design and probe design. Here, we used amplification-free, CRISPR-Cas13-based RNA detection for the identification of hantavirus genomic RNA in infection samples. We used the Cas13a ortholog from *Leptotrichia buccalis* (LbuCas13a) since we and others previously have shown that this ortholog has the highest sensitivity and activity compared to other Cas13 orthologs (15, 17).

Using Activity-informed Design with All-inclusive Patrolling of Targets (ADAPT), a system for the automated design of crRNAs (18), we designed crRNAs against the vRNA of the TULV or ANDV small (S) segment, which is the most abundantly expressed segment during hantavirus infection. We picked the top three ADAPT-designed crRNAs (named A, B and C) based on highest ADAPT-predicted activity. We then tested these three crRNAs individually, in pairwise combinations, or all together against *in vitro* transcribed (IVT) full-length S segment ANDV or TULV vRNA.

For TULV, we observed that when used individually, all three crRNAs yielded a signal at least 3 standard deviations above the water control when 10^8^ copies/μl of IVT target RNA were present in the sample. Unless defined otherwise, we used this cut-off of 3 standard deviations to define whether our assay had produced a statistically significant increase relative to the negative control. TULV crRNA A showed the highest fluorescent signal at 10^8^ copies/µl, followed by TULV crRNAs B and C (Figure 2A and B). Combining two crRNAs increased the signal compared to the individual TULV crRNAs, allowing detection down to ∼10^7^ copies/μl of the IVT target RNA when technical duplicates were analyzed (Figure 2A and B). This detection signal above background was higher for all pairwise combinations of crRNAs compared to the individual crRNAs. Furthermore, combining three TULV crRNAs resulted in higher and a less variable signal (Figure 2A and B). Thus, by combining all three crRNAs when detecting TULV vRNA, LbuCas13a can achieve the best performance.

**Figure 2.**
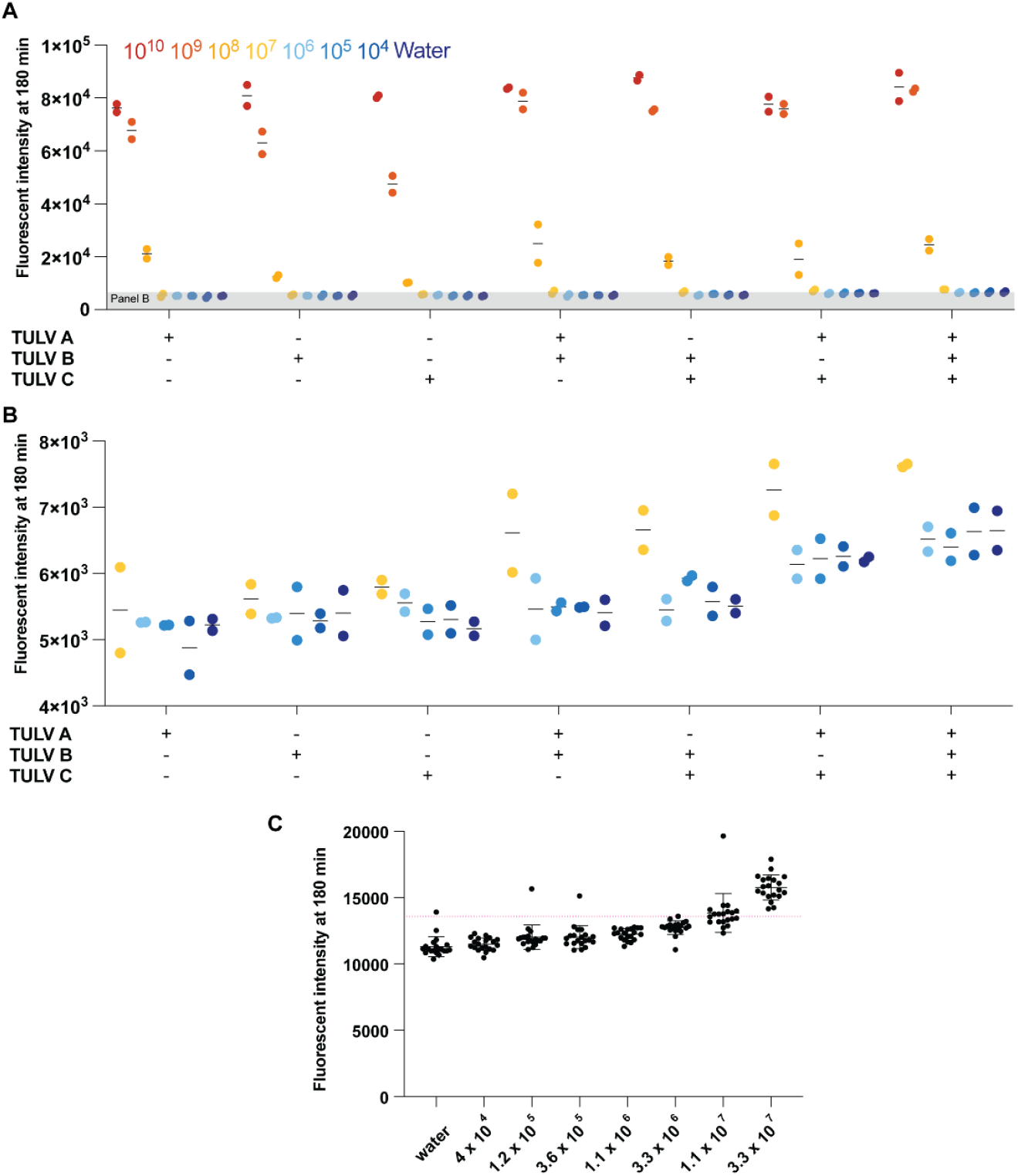
Validation of multi-crRNA Cas13-based amplification-free detection of TULV S-segment vRNA using IVT S-segment vRNA. **(A)** Detection of a 10-fold dilution series of *in vitro* transcribed v-sense S-segment TULV RNA using amplification-free Cas13-based approach using one, two, or three crRNAs. (B) Shows a subset of data from A. Each point represents one technical replicate. There are two technical replicates. (C) 3-fold dilution series to determine the limit of detection. The red line indicates our limit of detection cutoff of 3 standard deviations above the water input control. Each point represents one technical replicate. There are 20 technical replicates. The mean and standard deviation is shown.

For diagnostic use, a limit of detection (LOD) analysis based on many technical replicates is critical. To determine the LOD for the triple crRNA set, we repeated the LbuCas13a assay 20 times using serial dilutions of different concentrations of synthetic target RNA as input. Starting at 3.3 × 10^7^ copies/μl of IVT target RNA, we generated a 3-fold dilution series and measured each concentration using 20 technical replicates. We determined the limit of detection, which we defined as the point at which 19 out of 20 technical replicates were above background, to be 3.3 × 10^7^ copies/μl IVT RNA (Figure 2C).

When we use the ANDV crRNAs, the individual crRNAs A and B yielded a significant signal above background at 10^8^ copies/μl sample, whereas crRNA C was limited to 10^9^ copies/μl sample (Figure 3A and B). Combining two ANDV crRNAs increased the signal, allowing for detection down to 10^7^ copies/μl of target RNA when we combined crRNAs A and B or B and C, with the combination of crRNAs A and B producing the best signal (Figure 3A and B). However, combining A and C did not improve target RNA detection. When we combined all three crRNAs, we did not see a further improvement in sensitivity, keeping the detection at 10^7^ copies/μl RNA per sample (Figure 3A and B). Taken together, we conclude that combining ANDV crRNAs A and B produced the highest signal above background. As with TULV, we next generated a 3-fold dilution series and used this series for 20 technical replicates to determine the LOD. We found that the LOD for crRNAs A and B was 3.3 × 10^7^ copies/μl target RNA for 19 out of 20 technical replicates (Figure 3C).

**Figure 3.**
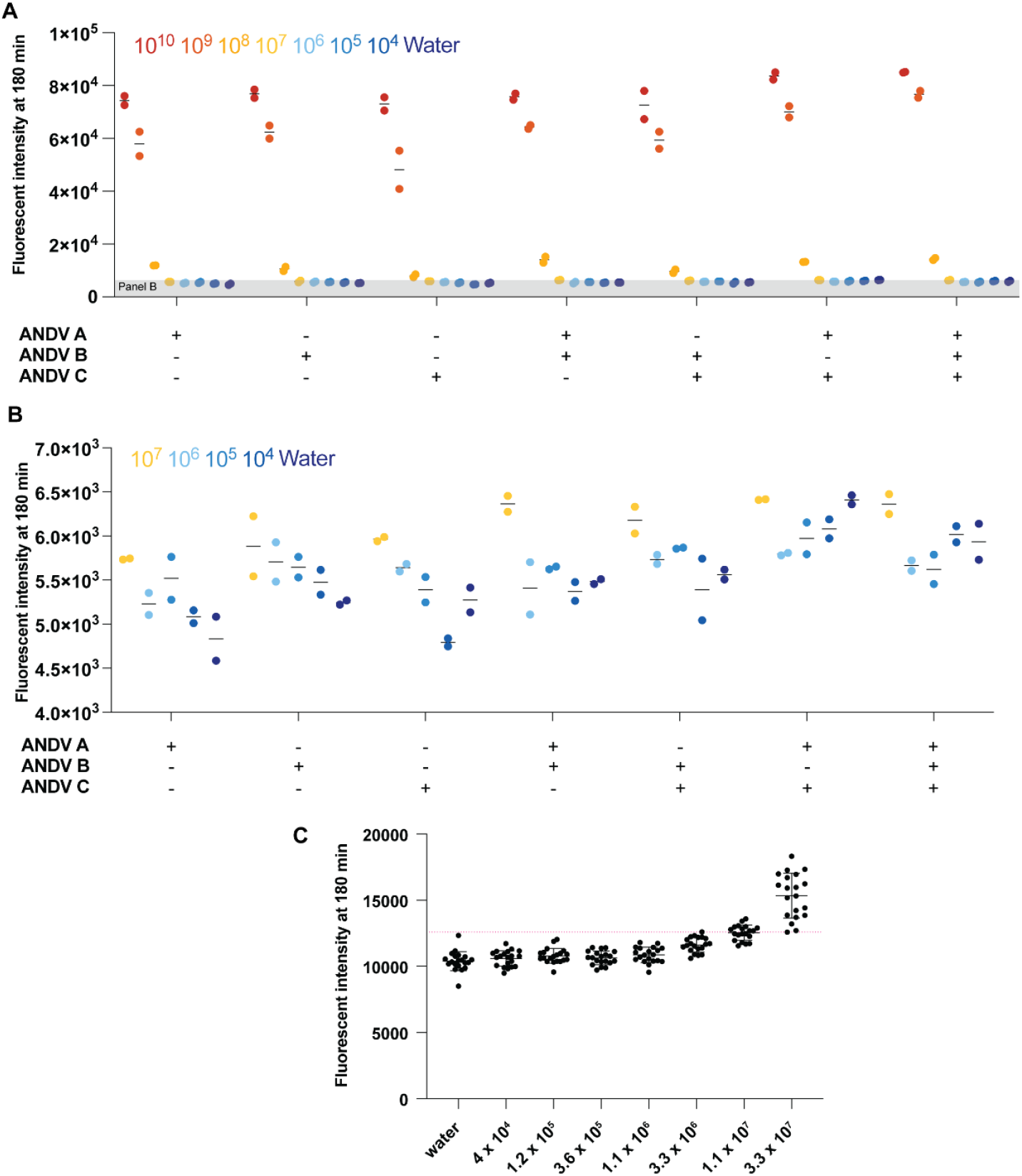
Validation of multi-crRNA Cas13-based amplification-free detection of ANDV s-segment vRNA using IVT s-segment vRNA. **(A)** Detection of a 10-fold dilution series of v-sense S-segment in vitro transcribed ANDV RNA using amplification-free Cas13-based approach using one, two, or three crRNAs. (B) Subset of data from A. Each point represents one technical replicate. There are two technical replicates. (C) 3-fold dilution series to determine the limit of detection. The red line indicates our Limit of Detection cut off, 3 standard deviations above the water background. Each point represents one technical replicate. There are 20 technical replicates. The mean and standard deviation is shown.

### Cas13-based detection of hantavirus RNA captures infection dynamics

After determining the LOD, we tested the ability of Cas13 to detect hantavirus vRNA in infected tissue culture cells. We infected human lung adenocarcinoma A549 cells, a typical cell line representing the primary site of hantavirus infection in humans, at an MOI of 0.1 with ANDV or TULV and extracted total RNA from cells at 6, 24, and 48 hours post infection (hpi). We then proceeded to detect S segment vRNA using Cas13 and RT-qPCR. ANDV infections were performed under biosafety level 3 (BSL-3) conditions.

Using Cas13, the ANDV S segment vRNA could be first detected at 24 hpi and its signal increased dramatically at 48 hpi (Figure 4A). The TULV S segment vRNA, however, was first detected at 6 hpi and its signal showed a more gradual increase towards 48 hpi (Figure 4B). Comparing these Cas13 results with the RT-qPCR readout, we found that the ANDV S segment RNA was detectable at 6 hpi by RT-qPCR (Figure 4C). In agreement with the Cas13 data, the vRNA signal increased dramatically at 24 and 48 hpi (Figure 4 A and C). The TULV vRNA was first detected at 6 hpi by RT-qPCR and increased steadily at 24 and 48 hpi, similar to what we observed using Cas13 detection (Figure 4 B and C). It is important to note here that the RT-qPCR was not specific to the vRNA molecules in the infection samples, since the RT step was performed using random hexamers. Thus, hantavirus cRNA and mRNA molecules had both contributed to the signal. This key difference between the Cas13 and RT-qPCR assays, combined with the amplification step in the qPCR assay, likely increased the apparent difference is sensitivity. To confirm that the differences in Cas13 and RT-qPCR signals were correlated with differences in ANDV and TULV growth kinetics, we performed western blots. We found a strong viral nucleoprotein (NP) expression at 48 hpi for ANDV, and a TULV NP signal at 24 hpi (Figure S1). Thus, the CRISPR-Cas13 method provides data that capture information on the viral infection cycle and that agree with traditional molecular methods.

**Figure 4.**
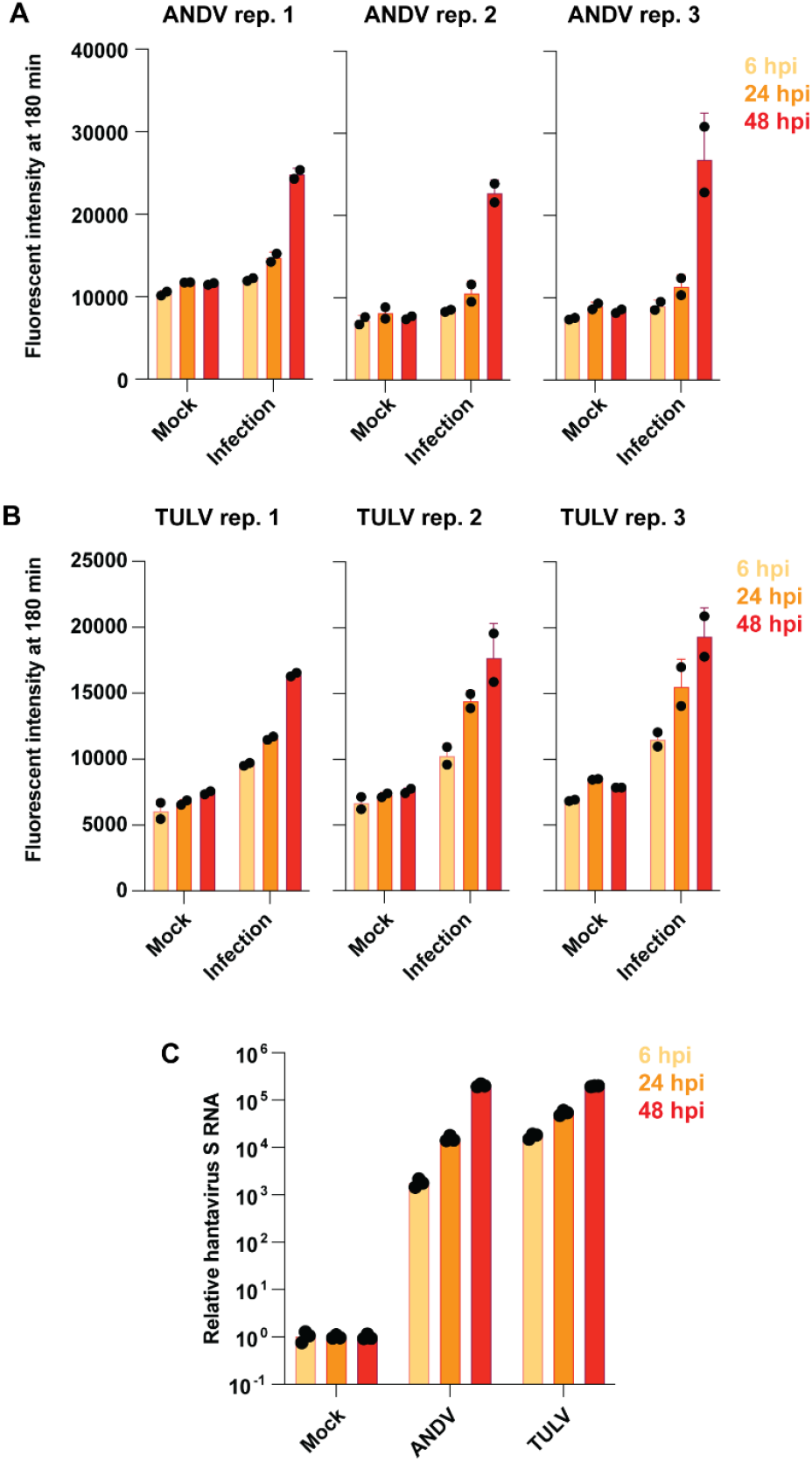
Comparison between multi-crRNA Cas13-based amplification-free detection of ANDV and TULV in tissue culture infected samples. Detection of ANDV or TULV S-segment in A549s infected with (A) ANDV or (B) TULV at MOI = 0.1 using multi-crRNA Cas13-based amplification-free detection or (C) RT-qPCR. Each graph depicted for Cas13 (A and B) shows one biological replicate with two technical replicates, represented as points. The graph for RT-qPCR (C) represents the mean relative fold change of S segment RNA compared to a housekeeping gene of three independent biological replicates.

### Combining crRNAs does not seem to improve IAV detection

Our above results show that using multiple crRNAs can increase the fluorescent signal and LOD of our LbuCas13a assay for detecting hantavirus RNA. To determine the general applicability of using multiple crRNAs for RNA detection, we tested whether it could be used for detecting IAV genomic RNA. To this end, we designed crRNAs targeting IAV vRNA or the replication intermediate, called the complementary RNA (cRNA), using ADAPT. We picked two crRNAs targeting the IAV vRNA (A and B) and three crRNAs (A, B and C) targeting the cRNA based on the highest ADAPT-predicted activity.

We first tested our set of crRNAs against a 10-fold dilution series of IVT segment 5 vRNA (Figure 5A). We found that IAV crRNA A supported a signal above background at 10^7^ vRNA copies/μl sample, while crRNA B only yielded a signal above background at 10^8^ copies/μl sample (Figure 5A). Combining the two crRNAs did not improve the overall detection of IAV vRNA (Figure 5A). We next tested our IAV crRNA set targeting the segment 5 cRNA against a 10-fold dilution series, starting at 10^10^ copies cRNA/μl sample (Figure 5B). All three crRNAs were able to produce a signal above background at 10^9^ copies/μl sample. When we combined two or three crRNAs, the assay yielded a signal above background at 10^8^ cRNA copies/μl sample (Figure 5B), indicating that combining crRNAs targeting the IAV cRNA does improve the detection.

**Figure 5.**
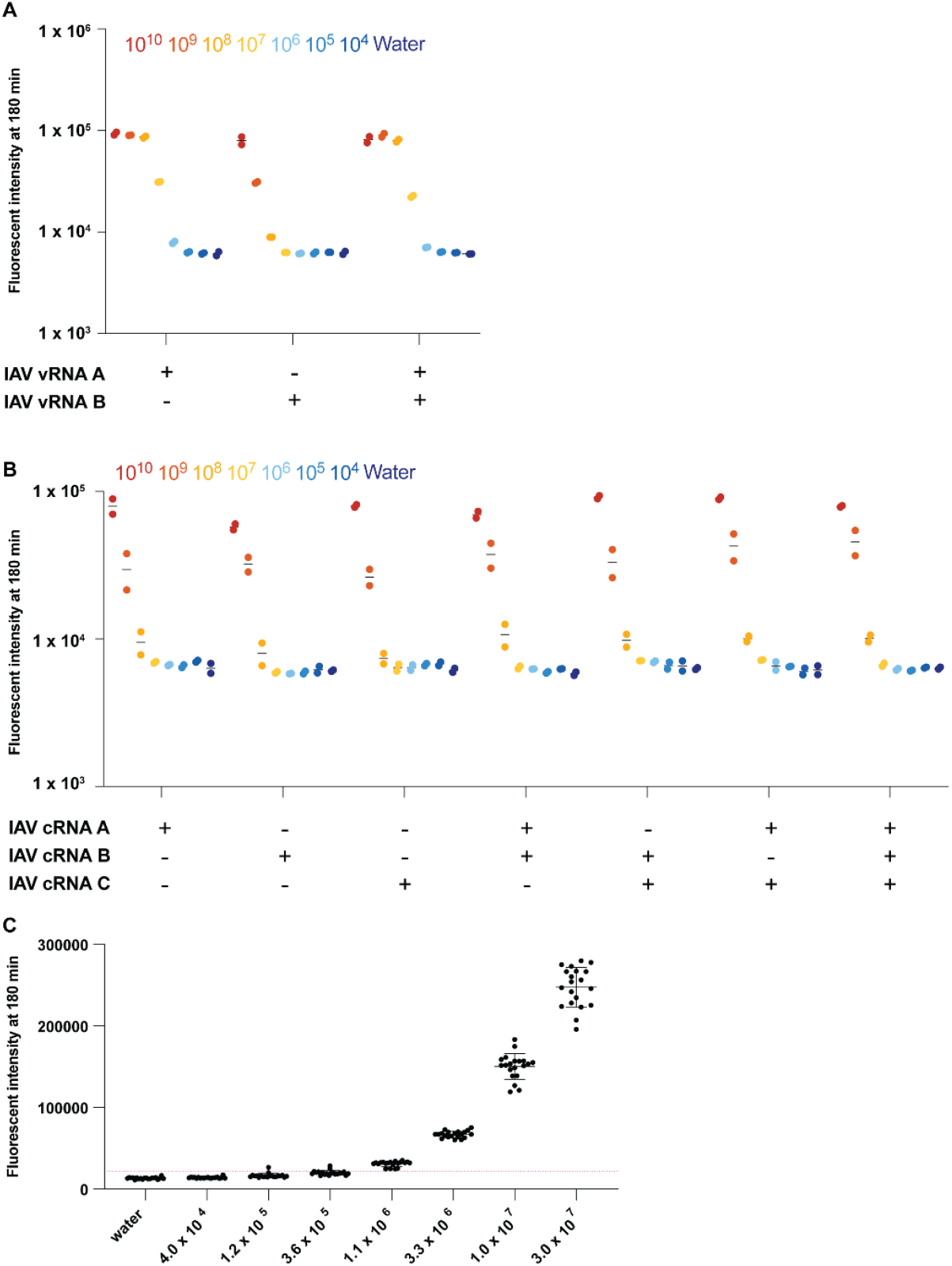
Validation of multi-crRNA Cas13-based amplification-free detection of v- and c-sense IAV segment 5 using IVT v- or c-sense segment 5 RNA. Detection of a 10-fold dilution series of (A) v-sense or (B) c-sense segment 5 IVT IAV RNA using amplification-free Cas13-based approach using one, two, or three crRNAs. Each point represents one technical replicate. There are two technical replicates. (C) 3-fold dilution series to determine the limit of detection. The dotted line indicates our LOD cut off, 3 standard deviations above the no input control. Each point represents one technical replicate (n = 20). The mean along with the standard deviation is shown.

We determined the LOD for the vRNA- and cRNA-targeting crRNAs using a 3-fold dilution series, starting at 3.0 × 10^7^ copies/μl, with 20 technical replicates at each concentration. We determined that the LOD of the vRNA-targeting crRNA A was 1.1 × 10^6^ RNA copies/μl of the IVT target RNA (Figure 5C). However, we were not able to detect a signal above the background for any combination of cRNA-targeting crRNAs and therefore did not determine an LOD for these crRNAs (Figure S2).

Finally, we tested whether our vRNA assay was specific to our strain of interest, WSN. We found that our assay was only able to detect A/Brisbane/59/07 (H1N1) and A/Oklahoma/05 (H3N2) at 10^10^ copies/μl sample, but not lower, and unable to detect A/Vietnam/1203/04 (H5N1) at all, indicating that the LOD for the correct IAV sequence is ∼3 orders of magnitude lower than the detection of incorrect IAV RNA and thus very specific (Figure S3).

### Cas13 can be used to detect IAV RNA in clinical samples

We next evaluated whether our IAV crRNAs could be used to diagnose clinical influenza samples. In addition to our vRNA-targeting crRNAs, we also tested the cRNA-targeting crRNAs. cRNA molecules are typically not present in IAV virions but are present in infected cells. We thus reasoned that we could use the vRNA- and cRNA-specific assays to get an indication of the amount viral and cellular material in the clinical samples. We had previously already shown that some 5S rRNA is present in IAV clinical nasopharyngeal swabs.

To test the clinical samples, we extracted total RNA from 20 clinical nasopharyngeal swab samples from patients that had tested positive for seasonal H1N1 or H3N2 using RT-qPCR. These samples included both male and female patients with ages ranging from 1 to 94 years of age (Table S2). We determined which samples were positive based on the fluorescence signal being 2 standard deviations above the mean of the no input (water) control samples (dashed line). Our vRNA-targeting assay detected IAV RNA in 15 out of 20 positive samples, i.e. 75%, and our cRNA-targeting assay detected IAV RNA in 17 out of 20 samples, i.e. 85% (Table S2, Figure 6). The cRNA signal was generally lower than the vRNA signal, suggesting that the nasopharyngeal swabs mostly contained viral and limited cellular material. Overall, our results demonstrate that our Cas13-based assay can be used to detect IAV RNA in clinical nasopharyngeal swabs.

**Figure 6.**
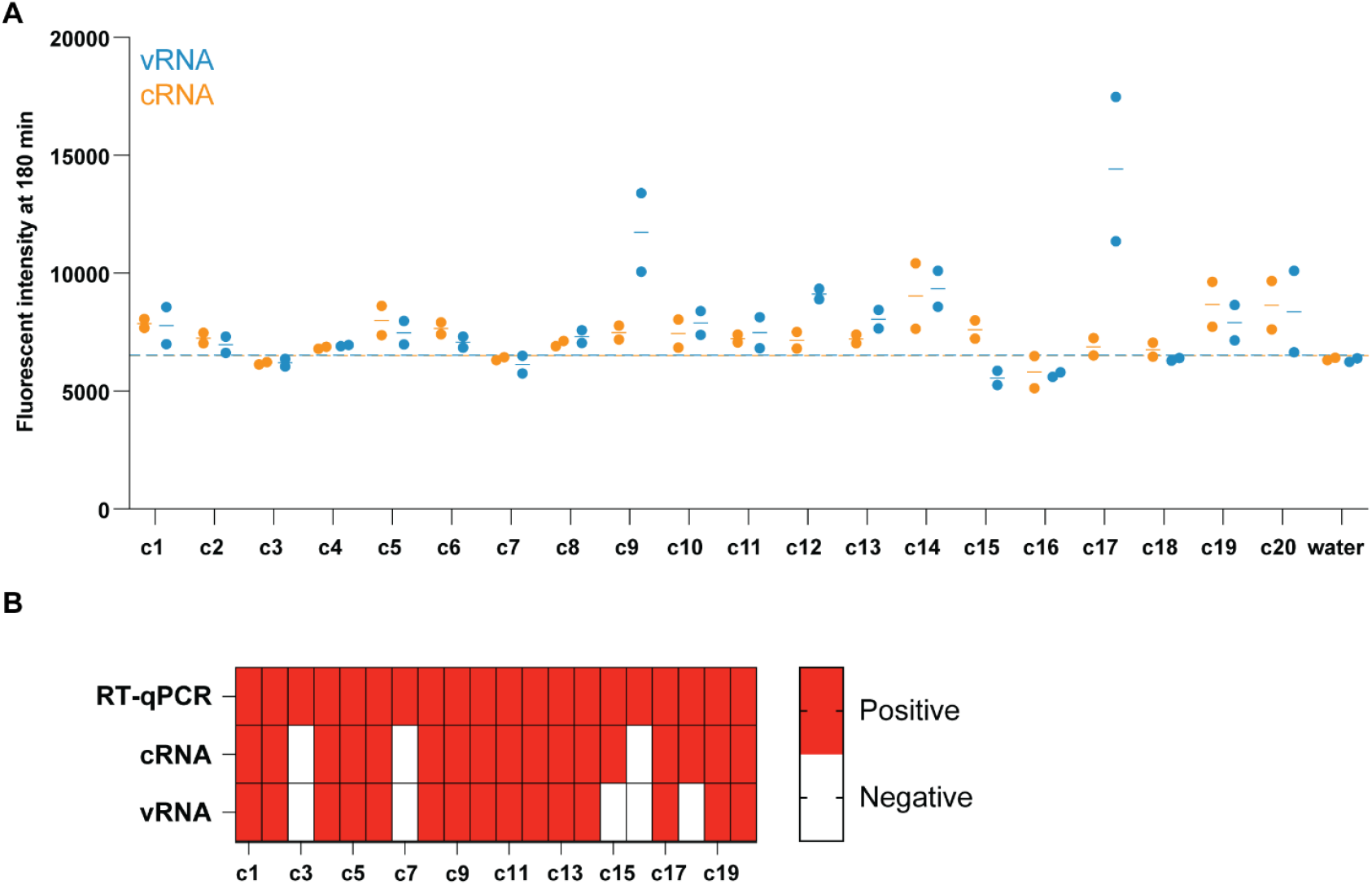
Cas13-based amplification-free detection of v- and c-sense IAV segment 5 in H1- or H3-RT-qPCR positive patient samples. (A) IAV segment 5 detection in RT-qPCR H1- or H3-positive patient samples (Table S2). The orange points represent fluorescent values obtained from the Cas13-based assay for cRNA detection, and blue represents fluorescent values obtained for vRNA detection. Dashed lines indicate cutoff for positive detection, determined to be 2 standard deviations above the no input (water) control. Each point represents one technical replicate. There are two technical replicates for each sample. (B) Comparison of the positive or negative diagnosis for each clinical sample based on the RT-qPCR data or the Cas13-based assays.

## Discussion

Here, we evaluated the effectiveness of an amplification-free, CRISPR-Cas13-based detection method for detecting RNA from zoonotic viruses such as hantaviruses and IAV. Our results demonstrate that combining multiple crRNAs can, in some cases, increase detection in agreement with previous SARS-CoV-2 data. However, we also observed that additional crRNAs do not always lead to improvement and that their impact depends on both the target and the crRNA sequences. Careful optimization of new CRISPR-Cas13 assay is therefore recommended.

For TULV and ANDV, combining two or three crRNAs enhanced the detection signal compared to single-crRNA conditions. Furthermore, we showed that this method can be used to track ANDV and TULV dynamics during infection in tissue culture and that the generated Cas13 data are consistent with RT-qPCR, meaning that Cas13, although less sensitive than RT-qPCR due to the lack of an amplification step, can still capture relevant trends.

In contrast to the hantavirus assay, the use of multiple crRNAs for IAV detection did not significantly enhance detection. For IVT RNA detection, a single v-sense crRNA was the most effective, outperforming an additional crRNAs even when combined. Similarly, the c-sense crRNAs showed minimal improvement in signal when used in combination, and the overall signal remained weak compared to v-sense detection. Even though a single crRNA provided the best Cas13 sensitivity for segment 5 vRNA, we were able to detect IAV RNA in clinical samples using both the v-sense and c-sense crRNAs.

In summary, while combining multiple crRNAs can improve detection for certain targets, such as hantaviruses, this approach does not translate to enhanced sensitivity across all viruses. Therefore, optimization must be tailored to the specific virus, segment, and crRNA set being tested. However, this optimization burden is relatively limited in the light of the simplicity of the assay and its potential as rapid diagnosis tool.

## Methods

### Preparation of synthetic RNA and primers

Primers were synthesized by Integrated DNA Technologies (IDT) and resuspended in nuclease-free water to 100 μM. Primers were stored at -20 ^°^C and were further diluted prior to analysis. crRNAs were synthesized by IDT and resuspended in nuclease-free water to 100 μM. crRNAs were stored at -80 ^°^C and were further diluted prior to analysis. Synthetic RNA targets were synthesized by IDT and resuspended in nuclease-free water to 100 μM.

### In Vitro Transcription of Target RNA

To generate IAV segment 5 RNA molecules, we first generated PCR products that added a T7 promoter to the 5′ terminus of the vRNA sequence. Specifically, we used a 2x Q5 mastermix (NEB), 100 ng of a pPolI-NP plasmid, and 1 µM of the primers T7min_FW and T7min_RV (Table S3). PCR products were analyzed by gel electrophoresis before purification using a PCR cleanup kit (NEB). T7 transcriptions were performed using 1 µg DNA template, 5 mM NTPs, 1 mM DTT, 1 U Murine RNase inhibitor (APExBIO), 1x T7 reaction buffer (Search Bioscience), and 100 U NxGen T7 enzyme. Reactions were incubated at 37 ^°^C for ∼3h, treated with DNase for 1 h, and then purified using a PCR clean-up kit (ZymoResearch). For transcription of the Hantavirus S segment, we used the same approach with Hantavirus-specific primers (Table S3).

### Cell culture

Human alveolar basal epithelial carcinoma (A549) cells (ATCC, CCL-185) and African green monkey kidney epithelial (Vero E6) cells (ATCC, CRL-1586) were commercially obtained. All cells were routinely screened for mycoplasma, and grown in Dulbecco’s modified Eagle medium (DMEM) containing pyruvate, high glucose, and L-glutamine (GeneDepot) supplemented with 10% fetal bovine serum (Gibco) and penicillin and streptomycin at 37 ^°^C and 5% CO_2_.

### Virus culture and titration

Andes virus (*Orthohantavirus andasense*) strain Chile-9717869 was a kind gift by Piet Maes (KU Leuven, Belgium). Tula virus (*Orthohantavirus tulaense*) strain Moravia/5302v/95 was a kind gift by Rainer Ulrich (Friedrich-Loeffler-Institute, Germany). ANDV and TULV were propagated in Vero E6 cells in DMEM supplemented with 2% fetal bovine serum, L-Glutamine, MEM Non-Essential Amino Acids and penicillin and streptomycin. During virus propagation, cell culture supernatants containing infectious virus particles were collected between 5 and 14 days post infection and replaced with fresh cell culture media. Pooled viral stocks were cleared from cellular debris by centrifugation (4,000 × g, 10 min, 4 °C) prior to three buffer exchanges in phosphate-buffered saline (PBS) and concentration using Amicon Ultra-15 centrifugal filter units (100-kDa molecular weight cut-off).

Concentrated viruses were titrated by immuno-plaque assays in Vero E6 cells. Briefly, Vero E6 cells were infected with sequential 10-fold dilutions of virus and overlayed with 1.2% Avicel in MEM containing 2% FBS, HEPES, MEM Non-Essential Amino Acids and penicillin and streptomycin. At 7 days post-infection, cells were fixed with 3.7% formaldehyde, permeabilized with 0.5% Triton X-100, blocked with 5% BSA in PBS and immunostained with cross-reactive primary monoclonal antibody targeting TULV NP (TULV1 (19)) and an IRDye-conjugated anti-mouse IgG (IRDye 800CW, 926-32210) secondary antibody. Near-infrared fluorescence signal of stained plaques was detected using an Odyssey CLx imaging system (LI-COR) and analyzed with the Image Studio software (LI-COR).

### Hantavirus infections

All work involving live ANDV was performed in the BSL-3 facility of the Hannover Medical School according to institutional biosafety requirements. For infections, approximately 2.5 × 10^5^ A549 cells were infected with TULV or ANDV at an MOI of 0.1 in DMEM supplemented with 2% fetal bovine serum, L-Glutamine, MEM Non-Essential Amino Acids and penicillin and streptomycin. At the indicated time points, cell culture supernatants were removed, and cells were washed once with PBS. Cells were lysed in TRIzol Reagent (Invitrogen) or RIPA buffer supplemented with 1% SDS.

### RNA Sample preparation

Influenza A virus clinical sample RNA extraction was carried out using Tri Reagent (Molecular Research Center, Inc) following the manufacturer’s instructions and as described previously (15, 20). Total RNA from hantavirus infected cells was extracted using TRIzol Reagent (Invitrogen) and the Direct-Zol RNA Miniprep Kit (Zymo Research) according to the manufacturers’ instructions. The RNA concentration was determined using a NanoDrop One spectrophotometer (Thermo Fisher) and diluted in RNase free water prior to analysis.

### RT-qPCR

Reverse transcription of RNA samples was performed using oligo(dT)/random hexamer primers and the PrimeScript RT Master Mix (TaKaRa Bio) according to the manufacturer’s instructions. Quantitative real-time PCR was performed using the TB Green Premix Ex Taq II (Tli RNase H Plus) Master Mix (TaKaRa Bio) on a LightCycler 480 Instrument II (Roche) according to the manufacturers’ instructions. Primers targeting human GAPDH (for: GAAGGTGAAGGTCGGAGTC; rev: GAAGATGGTGATGGGATTTC) and previously described universal (21) S segment hantavirus primers (for: CAGGAYATGVGRAAYACVATHATGGC; rev: CTCWGGRTCCATRTCATCMCC) were used. Delta-delta-cycle threshold (ΔΔCT) values were determined relative to the housekeeping gene GAPDH and normalized to the mean of the uninfected control group. Error bars indicate the standard deviations from three biological replicates (n = 3).

### Western blot

For western blot analysis, whole-cell lysates were obtained through cell lysis in RIPA buffer containing 1% SDS and sonication. Lysates were analyzed by SDS-PAGE and transferred onto nitrocellulose membranes. Membranes were stained with an Actin antibody (Invitrogen; MA5-11869) and a previously described Puumala virus NP antibody (clone 5E11) that is cross-reactive against ANDV and TULV NP (a kind gift by Indrė Kučinskaitė-Kodzė, Vilnius University). Primary antibody staining was detected using IRDye-conjugated secondary antibodies (IRDye 680RD, 926-68070; IRDye 800CW, 926-32210). Near-infrared fluorescent signals were detected using an Odyssey CLx imaging system (LI-COR) and analyzed with the Image Studio software (LI-COR).

### Cas13-based detection reactions

Each reaction contained 10 nM LbuCas13a, 4 mM HEPES pH 8.0, 12 mM KCl and 1% PEG, 2U/μL RNAse inhibitor murine (New England Biolabs), 0.25 μM 6UFAM (Table S3), 14mM MgOAc, 5nM crRNA (Table S3), and the reported amount of target RNA. First, LbuCas13a was combined with crRNA at a ratio of 2:1. The crRNA-Lbucas13a complex was subsequently added to a master mix containing HEPES, KCl, PEG, RNAse inhibitor, and MgOAc. For a final volume of 44 μL in 96-well plates. After mixing, 20 μL was transferred in duplicate to 384-well plates. The plate was then sealed and placed in a BioTek Synergy H1 Plate Reader and incubated at 37°C for 3 h. Fluorescence was measured every 5 min.

### Clinical samples

As previously described (15), nasopharyngeal samples were taken during routine testing from patients hospitalized at Addenbrookes Hospital during the 2016 – 2017, 2017 – 2018, 2018 – 2019 and 2019 – 2020 flu seasons. The study protocol was reviewed and approved by the Health Research Authority (IRAS ID 258438; REC reference 19/EE/0049). Patients were positive for either H1N1 or H3N2, but not other respiratory viruses, and samples were taken from a range of pathologies (asymptomatic to death). Per sample (typically 1.5 mL), 250 µl was used for total RNA extraction using Tri Reagent. RNA was dissolved in 10 µl RNase free water and stored at - 70 °C prior to Cas13 analysis.

### Statistical Testing

GraphPad Prism10 software was used for statistical testing. Unless otherwise stated, error bars represent standard deviation and individual data points indicate biological repeats.

## Acknowledgements

The authors thank Dr Hamid Jalal for support, and members of the Myhrvold and te Velthuis labs for useful discussions and suggestions.

